# Stretch-induced muscle responses under threat vs safety

**DOI:** 10.64898/2026.08.25.746965

**Authors:** Yvonne F. Visser, Bob Bramson, W. Pieter Medendorp, Karin Roelofs, Luc P.J. Selen

## Abstract

When in a stressful situation, making fast and accurate decisions is crucial. Previous work has shown that sensorimotor decisions can improve under threat. However, it is unclear if these improvements are achieved by improvements in perceptual or motor performance. Here, we present two hypotheses for how threat might influence motor preparation and use muscular stretch reflexes to test both. The task-unspecific hypothesis predicts that threat promotes motor preparation irrespective of the reach target, through tonic upregulation of the short latency stretch reflex. In contrast, the task-specific hypothesis predicts that threat increases sensory processing for a specific reach target, leading to direction-selective up- and down-regulation of the long latency stretch reflex. Participants were asked to reach to one of two targets that appeared shortly before a perturbation eliciting a stretch reflex, they performed this task either under threat of an electric shock or under safe circumstances. Skin conductance and heart rate results show that the threat manipulation significantly increased sympathetic activation, but not parasympathetic activation. Supporting the task-specific hypothesis, the EMG findings demonstrate a direction-selective modulation of the long-latency response of stretch reflexes, starting ∼100 ms after perturbation onset. Our results suggest that stress affects action preparation through upregulation of cortical visuomotor circuits.

## 1. Introduction

Freezing is a common physiological response in a threatening situation and is associated with hypervigilance, decreased heart rate and immobility (Blanchard & Blanchard, 1988; Blanchard et al., 1968; Carrive, 2006; Kozlowska et al., 2015). Though freezing may seem counterproductive, it appears to function as an opportunity for improved perceptual processing and preparation for an upcoming action (Livermore et al., 2021; Roelofs & Dayan, 2022). Indeed, perceptual sensitivity, orientation discrimination and perception of looming stimuli have all been shown to improve under threat (Lojowska et al., 2015; Vagnoni et al., 2012), which was closely correlated to threat-induced bradycardia (de Voogd et al., 2022; Klaassen et al., 2024; Lojowska et al., 2015). In addition, people become faster for accurate responses (Hashemi et al., 2019).

There is abundant evidence implicating the involvement of the amygdala and the periaqueductal grey (PAG), particularly the ventrolateral part (vlPAG), in the freezing response (Carrive, 1993; Hermans et al., 2013; Kozlowska et al., 2015; Roelofs & Dayan, 2022; Schipper et al., 2019). Both the amygdala and the PAG interact with visual cortical areas during threat-processing (Lojowska et al., 2018), which could explain the enhancing effects of threat on visual processing and perceptual decision making. Freezing-related heart rate deceleration has also been related to motor preparation and action planning (Gladwin et al., 2016). Indeed, there are direct connections from the vlPAG to brainstem areas and the spinal cord, which may facilitate this functional purpose (Castiglioni et al., 1978; Mantyh, 1983; Mouton & Holstege, 1994; Sillery, 2005; Tovote et al., 2016).

While previous studies have mainly focused on the perceptual consequences of freezing, here we examine how freezing could affect the preparatory motor state. We consider two hypotheses. First, freezing may directly affect action preparation due to increased arousal, thereby promoting the readiness to respond in the spinal loop. We refer to this as the ‘task-unspecific hypothesis’. Second, as freezing affects perception and there is a continuous flow between perception and action (Cisek, 2007), it could also modulate reflexes through cortical visuomotor loops. We refer to this possibility as the ‘task-specific hypothesis’.

To differentiate between these hypotheses, we relied on muscular reflexes elicited by a sudden stretch. The reflexive response to this stretch consists of different epochs. The so-called short-latency response (SLR, up to 45 ms after stretch onset) involves a monosynaptic spinal reflex loop and has a fixed size unless extensive training is undergone (Kurtzer, 2014; Scott, 2012; Wolpaw, 2010). The long-latency response (LLR, from 45 to 105 ms after stretch onset), involves a slower and more amendable (sub-) cortical loop. It can be modulated in a task-specific manner, either by the ongoing movement or future required movement directions, with stronger stretch induced responses in muscles that move the joint in the desired direction (Mutha et al., 2008; Reschechtko & Pruszynski, 2020). The timing of these reflex responses has been extensively reported and replicated, we used the same timings relative to stretch onset as described previously (Kurtzer, 2014; Mutha et al., Reschechtko & Pruszynski, 2020; Scott, 2012; Villamar et al., 2023; Wolpaw, 2010; 2008; Yang et al., 2011).

Figure 1 schematically outlines the two hypotheses, the presumed neural correlates and the expected effects on reflex activity. While the direct effect of freezing and the associated upregulation of PAG (fig. 1a) on spinal stretch reflex loops remains to be shown, there are clear indications of PAG modulatory influences on other spinal reflexes. For example, vlPAG stimulation in rats reduces withdrawal reflexes (Leith et al., 2010) and increases the H-reflex - the electrically-induced equivalent of the stretch reflex (Koutsikou et al., 2014; Koutsikou et al., 2015). In humans, inhibition of the main neurotransmitter of the sympathetic nervous system – noradrenaline – reduces the H-reflex (Palmeri et al., 1999). If similar influences from the vlPAG on the spinal cord exist, the task-unspecific hypothesis predicts a spinal-based increase of the reflexes in both the SLR and LLR epoch, irrespective of the required reaching direction (fig. 1a, bottom panel).

**Figure 1.**
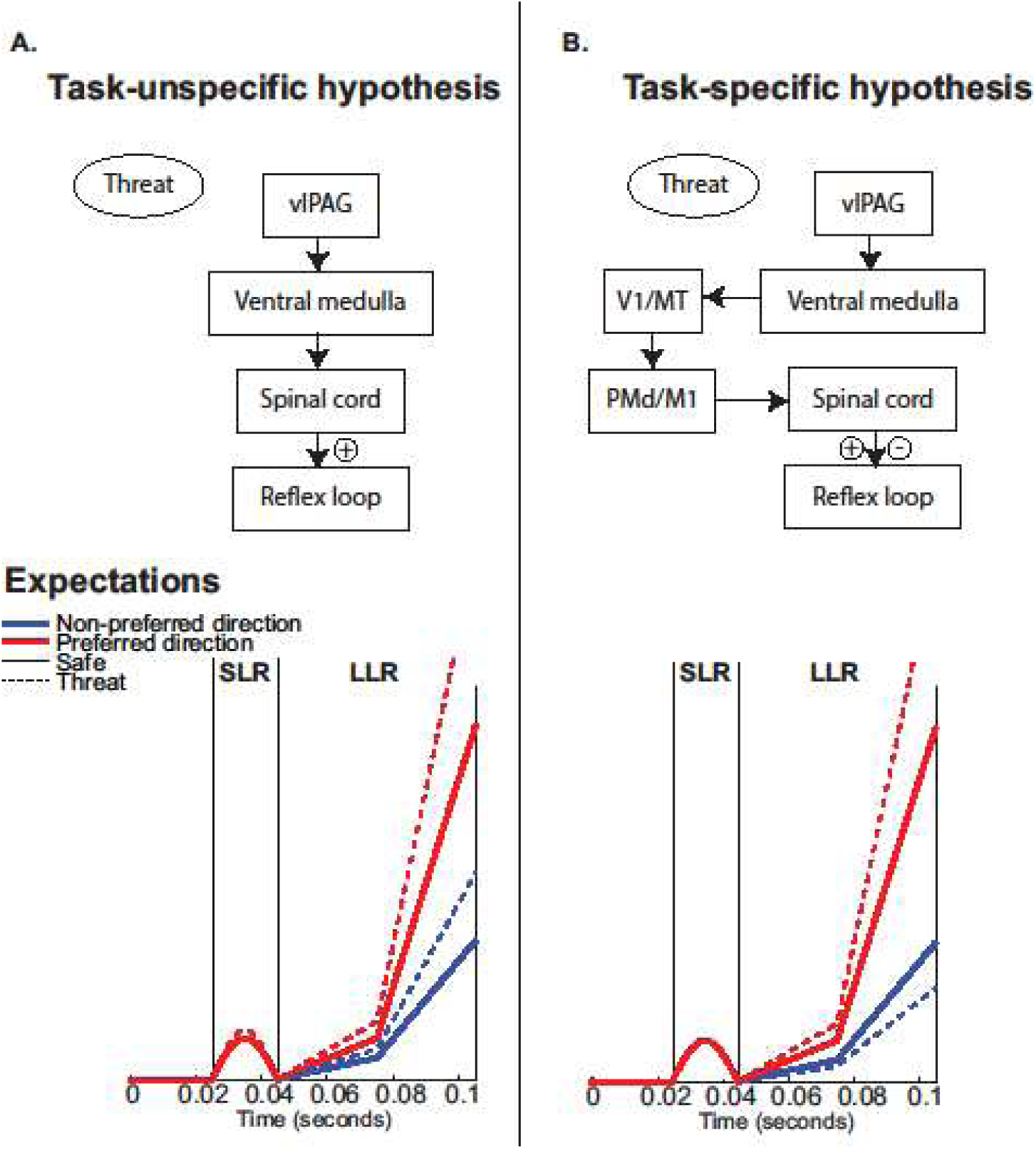
The two proposed hypotheses, the presumed neural correlates and the expected effects on reflex activity. **A)** If the threat-induced increase of vlPAG activity directly upregulates the neurons in the reflex loop, we expect an overall increase in reflex gains under threat (+). This increase is expected in all epochs of the reflex, including the SLR, and for reaches both in the muscle-preferred and non-preferred direction. **B)** If threat improves perceptual processing in visual areas, the information flowing to motor areas becomes more accurate and motor preparation can be tuned to the required response. This would result in an increase in reflex activity for the muscle-preferred direction (+) but a decrease in reflex activity for the non-preferred direction (-). We expect this effect to be visible in the LLR only, as it involves a cortical loop.

In contrast, the task-specific hypothesis (fig. 1b) implies that perceptual decision making is modulated by the vlPAG via a cortical relay, and that improvements are based on the continuous flow of information between sensory and motor execution areas (Cisek, 2007). As perception improves, evidence accumulates faster for the required action which results in a faster build-up of action specific long latency stretch reflexes (Selen et al., 2012; Visser et al., 2023). Thus, the task-specific hypothesis implies that the required action can be more rapidly prepared under threat, while alternative motor plans are inhibited. Hence, threat is expected to affect the LLR in a task dependent manner, whereas the SLR remains unaffected as it cannot be modulated in a task dependent manner.

In the present study, we elicited stretch reflexes while participants were preparing a reach to one of two possible targets, engaging different muscles. In some blocks, electric shocks could be delivered to induce threat, whereas other blocks were completely safe. Results show that stress modulates the later long-latency reflex response in a direction-specific manner, supporting the task-specific hypothesis.

## 2. Materials & Methods

### 2.1 Participants

Twenty-four healthy participants (17 female, 7 male, aged 18-61, mean 24.8 yrs.) took part in the experiment. The sample size was determined based on earlier studies on reflex modulation (Yang et al.,2011) and perception during freezing (Hashemi et al., 2019; Gladwin et al., 2016; Lojowska et al., 2015), using within subject designs.

All participants were self-reported right-handed and did not report any motor deficits, cardiovascular or endocrine disorders. Participants signed up through a local recruitment platform, gave informed consent before participating and were reimbursed for their time (€15/hour). The experimental procedures were reviewed by an independent local ethics committee – which had no formal objections - and complied with the declaration of Helsinki.

### 2.2 Set-up

#### 2.2.1 Robotic manipulandum

Participants were seated in front of a robotic manipulandum (3BOT, constrained to two dimensions to mimic the function of a vBOT (Howard et al., 2009), see fig. 2b), holding the robotic handle with their right hand. An airsled, positioned on a table below the handle, supported their forearm to facilitate frictionless planar movements. The robotic manipulandum, which could apply forces to the participant’s hand, tracked the handle’s position at 1000 Hz. A mirror was mounted above the movement plane, which reflected visual information (i.e., cursor position, targets, performance feedback) projected from a down-facing 27-inch screen (Asus MG279Q, 2560 x 1440 pixels, refresh rate: 144 Hz).

**Figure 2.**
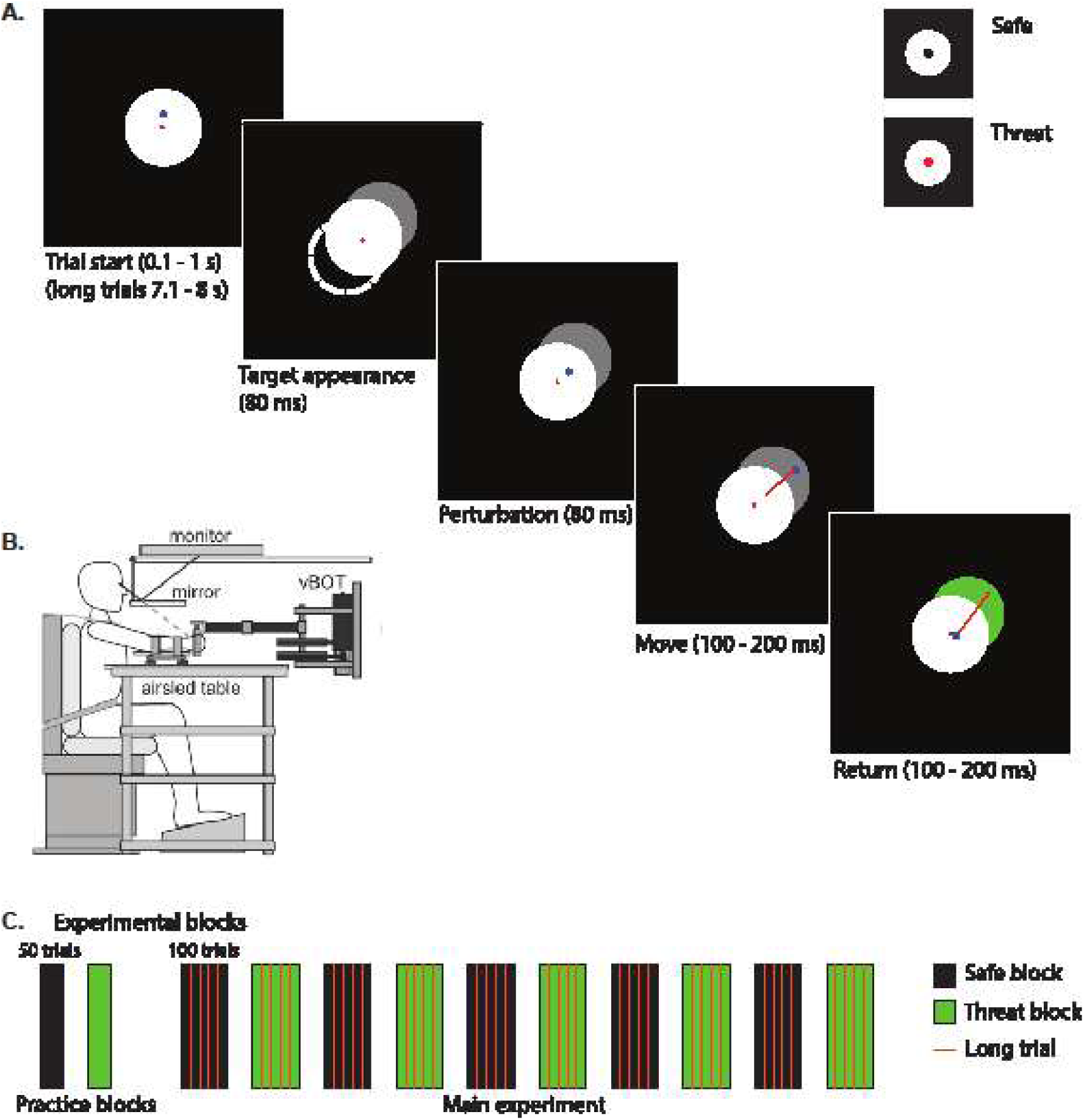
Experimental paradigm. **A)** Trial structure. Participants brought the handle of the robotic manipulandum into the center of the home position. After a variable delay of 0.1 to 1 second (or 7.1 to 8 s in the longer trials), a target appeared in elbow flexion or extension direction. The second target (dashed lines) is shown here for completeness, but was not visible to participants. After 80 ms of target presence, the hand was perturbed with an 80 ms, 1cm displacement in the extension direction, causing a stretch of the biceps. The perturbation was the ‘go’-cue for the participant to quickly move into the target area, which turned green once the handle had been in the target area for 100 ms. The participant then returned to the home position for the next trial. A central dot in the home position indicated whether the trial was ‘safe’ (black) or ‘threat’ (red). **B)** vBOT experimental set-up. Participants held the handle with their right hand while their forearm was supported by an airsled. Visual information was presented on the monitor and reflected by a mirror, such that the participant viewed them in the movement plane. **C)** Trial order. Participants performed two practice blocks of 50 trials, after which they completed ten main blocks of 100 trials, alternating between the safe (in black) and threat (in green) condition. Long trials were evenly spaced throughout the main blocks (orange lines).

#### 2.2.2 Electrophysiological measures

We recorded electromyography (EMG) of the right biceps brachii and triceps brachii muscles. The skin was scrubbed and cleaned before a wireless Delsys Trigno Research+ differential sensor was attached to the skin over the muscle belly of each muscle. A third EMG sensor was used to record an “electrocardiogram” (ECG) to determine the heart rate. We placed it on the left of the chest, below the clavicula, and confirmed sensor position based on visual inspection of the signal displaying a clear heartbeat. The system recorded both EMG and ECG data at 1925 Hz. Delsys Trigno sensors contain two reference sensors as well as two EMG sensors, so no separate reference electrode was placed. We acknowledge that these EMG sensors are not designed for ECG measurements, but initial piloting returned high signal quality and easily detectable heartbeats.

A GSR sensor (Brainvision) measured skin conductance at 500 Hz. The electrodes were attached to the proximal phalanges of the ring and little finger of the left hand and signal quality was confirmed with a Valsalva maneuver (*Practical tips for the GSR measurement*, n.d.).

EMG/ECG signals were time-aligned to the data from the robotic manipulandum by sending a trigger signal from the manipulandum software into the Delsys system at the time of stretch perturbation onset, while GSR signals were aligned through a photodiode connected to the Brainvision system.

#### 2.2.3. Threat-of-shock

To induce threat-of-shock, we administered aversive electric shocks to the participant. Stimulation electrodes were attached to the inside of the distal phalanges of the middle and index finger of the left hand. Due to a technical malfunction, we administered the shocks using an STM-ISOLA (BIOPAC) for the first 13 participants and an Isolated Bipolar Constant Current Stimulator (Digitimer DS5) for the last 11 participants. Shocks were randomly administered and were not related to task performance. Trials in which a shock was presented were excluded from the EMG and ECG analysis.

Prior to the experiment, the intensity of the shock was calibrated per participant to be perceived as unpleasant but not painful, using a standardized 5-step shock work-up (de Voogd et al., 2018; Klaassen et al., 2024). The shock current varied between 0.02 and 0.75 mA depending on the participant and had a fixed duration of 200 ms. For each participant, shock intensity remained constant throughout the experiment. Shocks were administered independently of task performance and could be presented at any time during a trial (except for the perturbation window, see below) with a 15% probability of occurring on any given trial in a threat block, though with a maximum of one shock per trial. A total number of 75 shocks were administered to every participant.

### 2.3 Paradigm

The paradigm served to elicit stretch reflexes for two opposing reaching directions when participants were and were not experiencing threat-of-shock. To this end, participants performed 1000 trials in a block-design. In threat blocks, an electric shock could be given at random times - irrespective of participants performance - while in safe blocks no shocks occurred. Importantly, shocks were performance-independent to induce an overall increase in arousal, without resulting in increased engagement in the threat blocks, which could have been a result of receiving punishment for errors in those blocks only. Blocks consisted of 100 trials and alternated between threat and safe, always starting with a safe block (fig. 2c). Safe and threat blocks were continuously cued by means of a coloured fixation point (black for safe, red for threat), and participants were allowed self-timed breaks between blocks.

Our task design was based on that used by Yang et al. (2011). They showed direction-specific modulation of stretch reflexes, with the perturbation starting 80ms after visual target presentation. That modulation increased with longer viewing durations. The short interval between target presentation and perturbation onset served for the information to travel through the nervous to modulate stretch reflexes in preparation of the required movement and avoiding visually evoked muscle responses. We aimed for a viewing duration in which direction-specific stretch reflex modulation occurred, but the tuning of the stretch reflex had not completely plateaued, such that any additional modulation by threat could still be shown.

A trial started when participants moved the cursor (veridically representing their hand position as a blue circle with a radius of 0.5 cm) into the home position (white circle with a radius of 7 cm presented in the center of the screen) and remained still (velocity < 1.5 cm/s). To indicate whether the participant was in a threat or safe block, a circular cue with a radius of 0.1 cm was presented in the center of the (white) home position. The cue was colored black in safe blocks and red in threat blocks (fig. 2a). After the participant had moved the handle into the home position, a variable delay between 0.1 and 1 s (drawn from an exponential distribution) preceded the appearance of a gray circular target with a radius of 6 cm in either the elbow flexion or extension direction. These reach directions were chosen to require either biceps or triceps engagement. The target circle partly overlapped with the home position, resulting in a half-moon shape with a maximum width of 4 cm. A mechanical perturbation started 80 ms after target appearance (which was verified with a photodiode), pushing the handle 1 cm in the elbow extension direction within 80 ms (corresponding to a velocity of 12.5 cm/s). This perturbation stretched the biceps muscle, allowing us to study the stretch induced changes in the muscle activations. Participants were instructed to reach toward the target as soon as they noticed the perturbation, to maximize preparation based on target appearance. Once the handle had been inside the target area for 100 ms, it turned green and the participant returned the handle to the home position for the next trial. If the handle speed exceeded 1.5 cm/s before perturbation onset or the participant failed to reach the target area within 1 s after perturbation onset, the trial was marked as missed and repeated right away. The original missed trial was not included in the analysis.

To study heart rate changes and skin conductance, we introduced 40 long trials (20 for safe and 20 for threat blocks) - equally spaced throughout the experiment (see fig. 2c). In these trials the target delay was between 7.1 and 8 s instead of 0.1 and 1s. The delay between target and perturbation onset remained at 80 ms, and EMG data from these trials was also included in the analyses.

A practice block of 100 trials (50 in the safe and 50 in the threat condition) was performed prior to the main experiment. The trials in this block were identical to the main experiment, except for the timing of the perturbation relative to target presentation. Instead of an 80ms viewing duration, the participant had seen the target for 200 ms before perturbation onset. Data from the practice block were not analysed.

### 2.4 Data analysis

#### 2.4.1 Behavioral analysis

To investigate possible improvements in performance from safe to threat trials, we compared several kinematic measures between the two conditions. First, movement time was quantified as the time between the end of the perturbation and the cursor entering the target area. Movement time was used as a proxy for movement vigor, which has previously been shown to decrease as accumulated evidence increased (Selen et al., 2012). Second, we counted the number of missed trials, i.e. trials in which the participant initiated a move before perturbation onset, or in which the reaching duration was above 1 s. Finally, to quantify possible differences in partial errors – where participants start their reach in the wrong direction - we compared the trajectory deviation from a straight line between the safe and threat condition at the beginning of the reach. To this end, we computed the accumulated lateral distance of the cursor relative to the line directly connecting the start location and center of the target up to the point that the cursor crossed the threshold of 1cm perpendicular to this line in the direction of the target. This analysis was designed to capture the distance from the ideal movement direction during the initial part of the reach (i.e. 1 cm away from the start). For a partial error, this distance should be relatively large, whereas it would be close to zero if the participant reaches straight to the target.

#### 2.4.2 Electrophysiological preprocessing

EMG data was rectified and low-pass filtered at 45 Hz (5^th^ order Butterworth bidirectional filter), before segmenting into trials from -0.5 to 1 s relative to perturbation onset. EMG epochs were further z-scored to facilitate averaging over participants. A per-trial baseline normalization was performed by dividing the preprocessed trial EMG by its average value just prior to perturbation onset (-0.5 – 0 s). We defined the two reflex epochs: the short latency reflex (SLR) from 25 to 45 ms and the long latency reflex (LLR) from 45-105 ms after perturbation onset. EMG activity from 105ms onwards was considered voluntary.

Heart rate and skin conductance were analyzed based on the long trials only. The ‘electro-cardiogram’, i.e. its proxy based on a differential EMG electrode, was segmented from -5 to + 10 s from trial start, i.e. the moment the cursor entered the home position. In this segmented ‘electro-cardiogram’ signal we determined the heartbeats. Since we used an EMG instead of an ECG montage, we could not use a standard Pan-Tompkins algorithm on the PQRST-complex. Instead, we first z-scored the signal and subsequently used Matlab’s ‘findpeaks’ function to determine peaks that exceed a z-value of 2 (i.e. 2 standard deviations greater than the mean). Note, the sign of the R-peak varied in each participant due to the placement of the EMG electrode. In participants where this peak was negative, we inverted the signal. To exclude spurious peaks, e.g. due to left pectoralis muscle contractions, we computed a rough estimate of the inter-beat interval and found for every peak the next peak that is closest to this inter-beat interval. All other, spurious, peaks are removed. Next, we computed the inter-beat interval and converted it into beats-per-minute (BPM). We assigned this BPM-value to the whole inter-beat interval. To quantify heart-rate deceleration we subtracted, per trial, the BPM value between -1 and 0 s relative to trial onset.

The skin conductance signal was lowpass filtered at 125 Hz and bandpass filtered at 50 Hz, before epoching at -5 to 10 seconds relative to long trial onset. We defined the skin conductance response (SCR) as the square root of its maximum value between 0.5 and 5.5 s after trial start, subtracting the value between -1 and 0 s relative to trial onset (see Boucsein et al. (2012) for conventions). The difference in average skin conductance during long trials of safe vs threat blocks (safe – threat) was computed for plotting purposes only.

Trials in which a shock was presented (7.5% of all trials) were excluded from analyses.

#### 2.4.3 Statistics

Paired-sample t-tests were used to test for behavioral differences in reaction time, movement duration, missed trials and lateral deviation. For the SCR between safe and threat, a paired t-test was used as well.

We used a Bonferroni-corrected 2-sided Wilcoxon signed rank test to evaluate differences in heart rate deceleration, segmenting the signal into windows of 0.5 s between - 7 and 0 s relative to target onset.

To investigate differences in reflex and voluntary EMG between safe and threat blocks, we compared EMG in both the biceps and triceps muscle for the two different reach directions. We performed separate cluster-based permutation tests (Maris & Oostenveld, 2007) for the task-unspecific and task-specific hypothesis predictions, testing for both an in-and decrease of the EMG activity in threat compared to safe for the non-preferred direction in each muscle (see fig. 1). We quantified the threshold for the test statistic in the clustering based on the t-value between safe and threat. Cluster-based permutation tests were performed on the EMG signal ranging from 0 to 0.5 s after perturbation onset.

Then, to improve power, we combined EMG measurements from both reach directions and both muscles. To this end, we subtracted average EMG activity in safe from that in threat trials for the four combinations of biceps/triceps and flexion/extension. Under the task-unspecific hypothesis, this subtraction should always result in a positive number: EMG activity is higher under threat than in safe conditions. However, under the task-unspecific hypothesis, the sign of the result depends on the preferred direction of the muscle. For example, for the biceps, EMG activity for targets in the elbow flexion direction should be higher for threat than for safe conditions, but activity of the biceps should be lower for threat than for safe conditions for targets that require elbow extension. This is due to the antagonistic function of the biceps in the case of an elbow extension. For the triceps, there is a similar expectation, but flipped across the reach directions, since it works as an agonist in the elbow extension direction. When we sum the muscle-specific subtractions based on their (non-) preferred direction, we get opposite predictions for each hypothesis. Under the task-unspecific hypothesis, we expect no interaction: the preferred and non-preferred direction should both show an increase for the threat condition compared to the safe condition. Under the task-specific hypothesis, however, we expect the EMG activity to be increased for the preferred, but decreased for the non-preferred direction when experiencing threat (fig. 5a). In the cluster-based permutation test, we computed t-values for safe vs threat in the preferred minus the non-preferred direction for the individual muscles and added them before determining the clusters. Subsequently we found the cluster with the largest summed t-value over a continuous time window with a t-value above zero.

## 3. Results

Participants were instructed to reach to a briefly (80 ms) visible target that required either elbow flexion or extension as soon as they felt a mechanical perturbation that elicited a stretch reflex. Half of the blocks were safe, whereas in the other half participants were under threat of an electric shock. We recorded EMG, heart rate and skin conductance and compared behavior and electrophysiological responses between the safe and threat blocks.

### 3.1 Skin conductance, heart rate & behavior under threat of shock

We used the SCR and heart rate deceleration to quantify freezing in the 40 long trials.

Figure 3a shows the time development of skin conductance for long trials in the threat compared to safe condition. When computing the peak value of the skin conductance on each trial (the SCR), we found an increase for threat compared to safe trials (fig. 3b, t(23) = 3.10, p = 0.005), indicating higher sympathetic nervous system activation in the threat than in the safe block. Heart rate showed an overall reduction relative to trial start, in the same time window as skin conductance increase, which is compatible with the notion of freezing-related bradycardia. Heart rate reduction showed no differential effect for safe and threat trials on a group level (fig. 3c, p>0.05).

**Figure 3.**
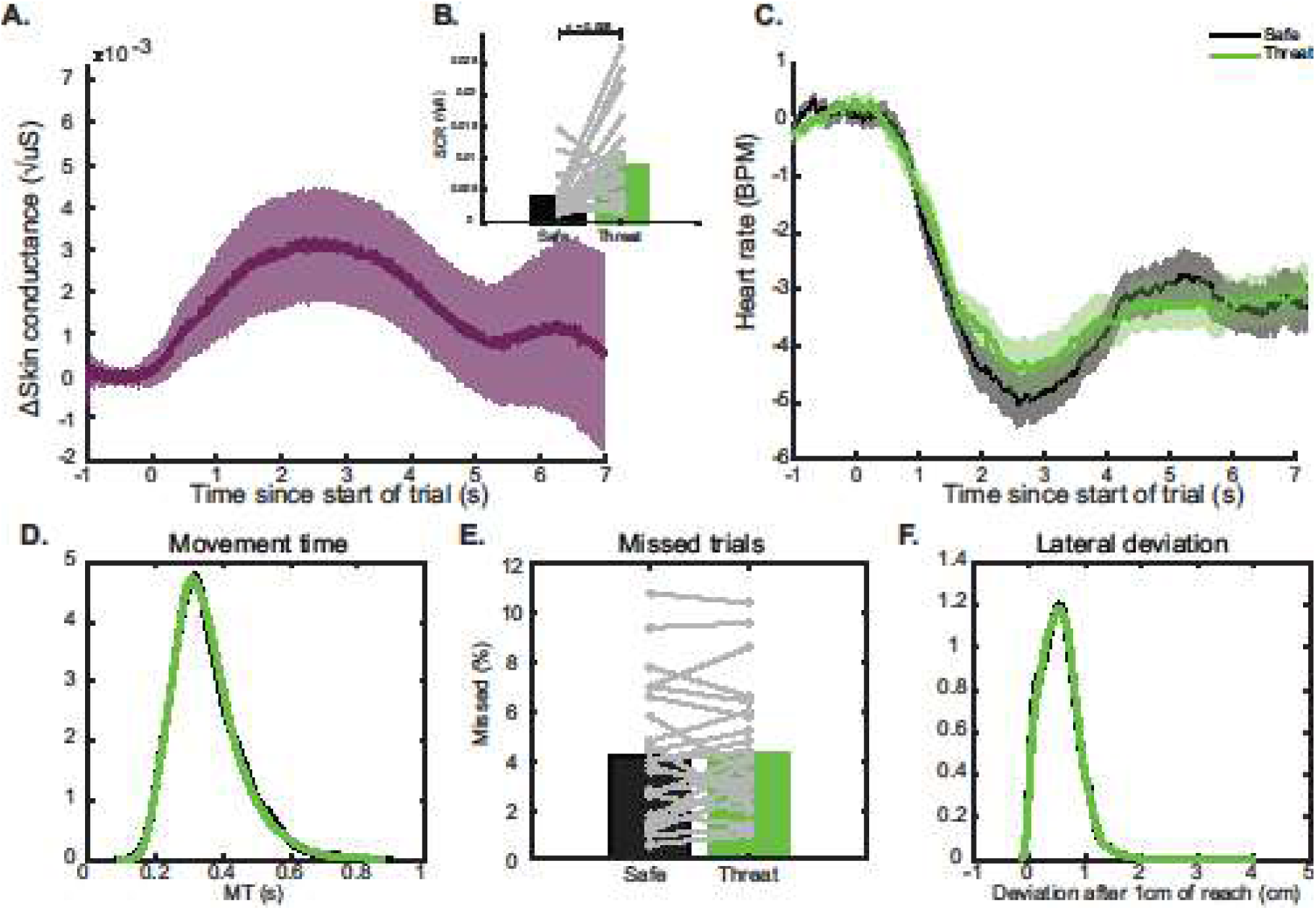
Skin conductance response, heart rate deceleration and behavior in threat and safe trials. **A)** Skin conductance in threat compared to safe blocks, relative to the start of a long trial. **B)** Skin conductance response (SCR), or the peak of the skin conductance between 0.5 and 5 seconds for each trial. SCR in threat trials was significantly higher than in safe trials. Gray open circles and lines represent individual participants. **C)** Heart rate deceleration relative to the start of a long trial, baselined to the period prior to trial start. We did not find a difference in heart rate deceleration between safe and threat blocks. **D)** Distributions of movement time: the time between the end of the perturbation and when the handle entered the target area. **E)** Percentage of trials that were missed due to a participant starting their movement before the perturbation or reaching too slowly. Gray open circles and lines represent individual participants. **F)** Distributions of trajectory deviation. None of the behavioral measures resulted in a significant difference between safe and threat blocks.

To check for changes in performance between safe and threat, we compared movement duration, percentage missed trials and lateral deviation from a straight reach trajectory between the threat and safe blocks. We found no significant differences in any of these measures (fig. 3def, p>0.05).

Since we did not find the classical signatures of threat-related processing nor perceptual improvement, we will speak of stress instead of threat for interpretation of the results.

### 3.2 Long latency reflexes and voluntary EMG are modulated by reach direction and different for the safe and threat condition

Figure 4 shows the average EMG activity across participants for biceps (panel A and C) and triceps (panel B and D), separated by reach direction and threat condition. Both in the LLR window (45-105ms) and the voluntary window (>105ms) a clear modulation of EMG activity with reach direction is observed, whereas for the SLR window (20-45ms), stretch induced activity seems unaffected by reach direction. Especially in the LLR and voluntary epoch, EMG activity is heightened for the preferred (when the muscle acts as an agonist) compared to the non-preferred (when the muscle acts as an antagonist) direction.

**Figure 4.**
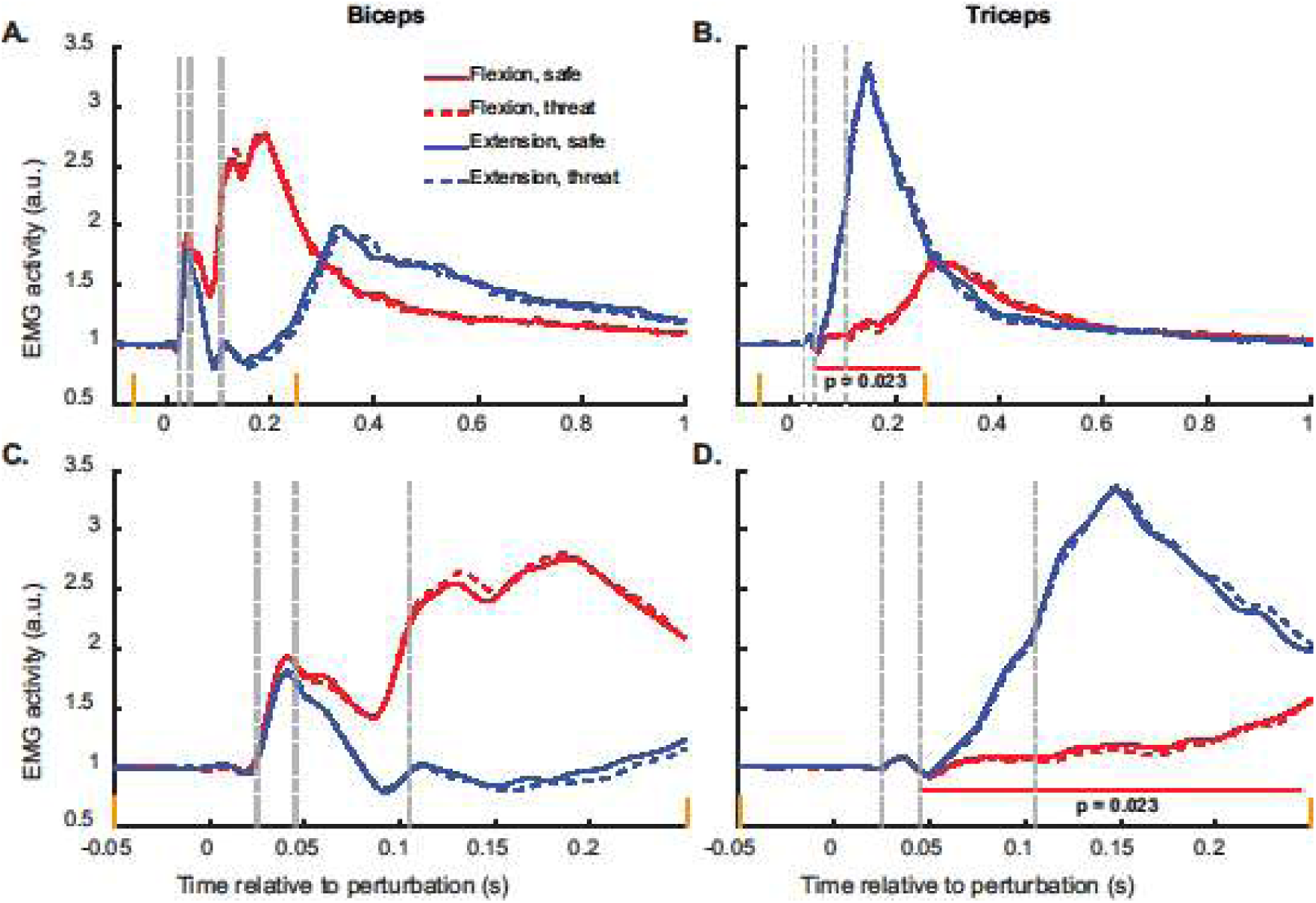
Grand average EMG. **A** and **B** Biceps and triceps average EMG activity across participants, for the SLR, LLR and voluntary epoch, separated for reach direction (red for flexion, blue for extension) and threat condition (solid for safe, dashed for threat). **C** and **D** same as A and B but zoomed in on the SLR, LLR and early voluntary window. Orange ticks indicate the same point on the x-axis for the overview and zoomed version of the plot. The vertical dashed lines indicate the SLR, LLR and the start of the voluntary window. A one-sided test for the perceptual hypothesis revealed a significant cluster between 0.0454 and 0.2443 s after perturbation for the triceps when reaching for the flexion target (red horizontal line in B&D).

EMG traces for safe (solid) and threat (dashed) blocks show a similar response for the same reach direction (fig. 4). According to the task-unspecific hypothesis, the EMG amplitude should be slightly higher for all muscles and all reach directions in the threat condition (fig. 1a), whereas according to the task-specific hypothesis the EMG responses should be more strongly modulated by reach direction in the threat compared to the safe condition. This should be observable as an increase in muscle activity for the preferred direction and a decrease in muscle activity for the non-preferred direction (fig. 1b). Some evidence for the latter-task-specific - hypothesis, is visible when comparing the dashed and solid lines in figure 4. In particular, the biceps EMG is lower under stress for extension and higher for flexion. The opposite pattern is seen in the triceps.

To statistically verify these observations, we performed four cluster-based permutation tests, one for each combination of reach direction and muscle, contrasting the safe and threat condition. This resulted in a significant (p = 0.023) cluster between 0.045 and 0.244s (LLR and voluntary window) for the triceps for its non-preferred (flexion) direction (fig. 4bd). The significant cluster shows lower activity for the threat condition than for the safe condition, which is in accordance with the task-specific but not the task-unspecific hypothesis (fig. 1). Exact effect sizes are unreliable for these types of tests (Lehongre, 2017), but the approximate cohen’s d suggests a small effect size. However, none of the other tests resulted in a significant cluster.

To increase statistical power, we combined trials from different reach directions and both muscles by subtracting EMG activity in the safe from the threat condition for both preferred and non-preferred reaching directions per muscle (see methods and fig. 5a). Then, we compared the resulting difference t-value timeseries for the preferred versus the non-preferred direction. Our two hypotheses give different predictions for those timeseries. According to the task-unspecific hypothesis, the amplitude of all reflexes should increase for threat, regardless of reach direction, so subtracting the timeseries for preferred from non-preferred direction should result in no difference (fig. 5a). For the task-specific hypothesis, however, the predicted differences in both reach directions are of opposite sign and should result in a remaining difference when subtracting the preferred from the non-preferred reach direction (fig. 5a). We used a cluster-based permutation test to look for differences between those threat-safe difference signals, collapsed over both reach directions and both muscles. This test, contrasting the preferred and non-preferred reach directions, resulted in a significant cluster (p = 0.017) in the early voluntary EMG (0.102-0.320 s), supporting the task-specific hypothesis (fig. 5b).

**Figure 5.**
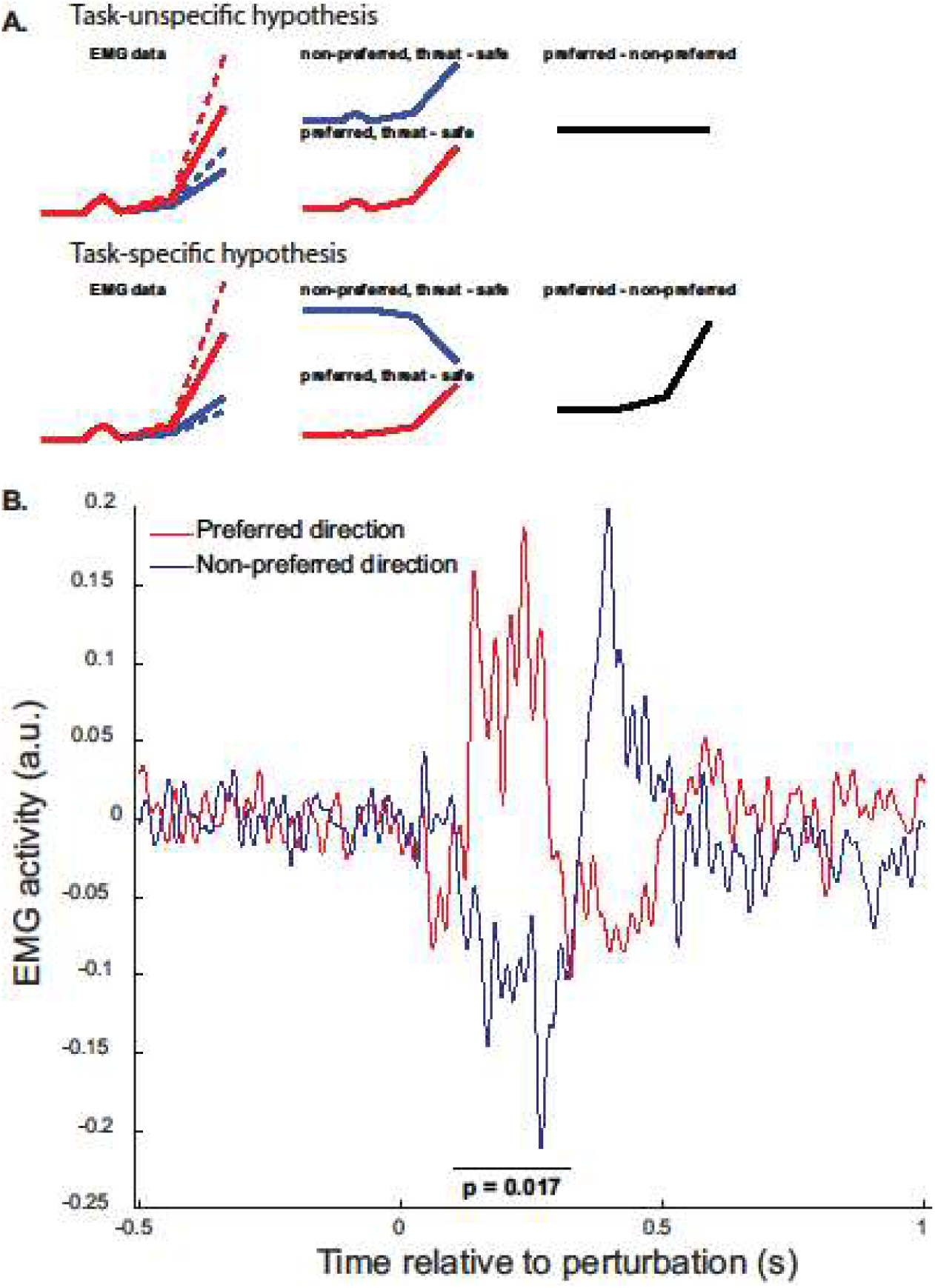
Contrasting overall EMG activity for the safe and threat condition. **A)** Expectations for the contrast between preferred and non-preferred direction in both muscles for the task-unspecific and task-specific hypotheses. Under the task-unspecific hypothesis, there is no interaction between reach direction and preferred direction and would mean the differences between the safe and threat condition are cancelled out when contrasting the directions. Under the task-specific hypothesis, a significant difference between the directions is expected. Since this prediction is in the same direction for both muscles, we can combine them to increase power. **B**) Subtraction of EMG activity in safe vs threat conditions for both biceps and triceps in their preferred and non-preferred reach direction. A significant cluster is found between 0.102 and 0.320 s after perturbation onset, where the contrast between safe and threat for the preferred direction is different to the contrast between safe and threat for the non-preferred direction. This result supports the task-specific hypothesis.

## 4. Discussion

We investigated how a threat of shock manipulation affects motor preparation, quantified as stretch reflex magnitude, by asking participants to reach for a target in either the elbow flexion or extension direction, as soon as they felt the stretch perturbation.

Participants performed this task in both a safe condition and under threat-of-shock. We recorded EMG, heart rate and skin conductance during the experiment. We show that stress modulates late reflexes in a way that is consistent with patterns that would be expected if the PAG modulates task-specific cortical visuomotor loops, rather than a task-unspecific general arousing influence on the reflex loop.

We tested two opposing hypotheses on how stress might bias action preparation: the task-unspecific hypothesis, which predicts general upregulation of both the SLR and LLR; and the task-specific hypothesis, which predicts stronger direction-modulation in the LLR under threat (fig. 1). When combining both reach directions and muscles, we found increased EMG activity under threat for the muscles’ preferred direction between 100 and 300 ms after perturbation onset, while activity was reduced under threat for the non-preferred direction (fig. 5b). The timing of these differences is relatively late, falling in the voluntary epoch of the EMG response to stretch. However, being this early in the voluntary epoch, we might still assume an effect of motor preparation on this activity. This result supports the task-specific hypothesis, i.e. that stress-induced improvements in perception lead to more specific motor preparation. Additionally, we found a significant decrease in EMG activity early in the long latency reflex for the triceps when reaching to the flexion target, which is also in line with the task-specific hypothesis (fig. 4b and d). The timing of those stress-effects is in line with previous results suggesting a continuous flow of information between perceptual and motor areas (Cisek, 2007; Donner et al., 2009; Gold & Shadlen, 2003; Michelet et al., 2010). This idea suggests that evidence accumulation does not occur only in sensory areas, but partial information is continuously sent further downstream. Though our results suggest that this information stream does not go as far as the spinal cord, it is likely that the effect in the LLR depends on some type of central sampling (perhaps of the motor cortex or the brainstem), for which there is enough time in the LLR but not the SLR. Finding the effect in the LLR strengthens the evidence for the task-specific hypothesis, since general upregulation as suggested by the task-unspecific hypothesis could have been accomplished by tonic increases in spinal cord excitability independent of perceptual processes in the central nervous system. Our findings align with a recent study in which the locus coeruleus was electrically stimulated to increase norepinephrine projections. The authors show that sensory processing is enhanced, as opposed to a simple change in response speed (Su et al, 2025).

Even though the differences in EMG activity between safe and stress align with the task-specific hypothesis, they occurred later than we anticipated (102-320 ms after perturbation). From 105 ms, EMG activity is generally classified as ‘voluntary’, not as a reflexive response to the stretch perturbation. This would suggest that the reflex itself is not altered by the threat condition, but only the voluntary muscle activity to perform the reach. However, the observed difference starts very early in this voluntary epoch, and could well reflect a transition from the reflex to voluntary activity. In fact, the dichotomy between reflexes as automatic and unchangeable and voluntary control as being cerebral and explicit has been questioned (Krakauer, 2019). For the triceps muscle, modulation was found soon after the perturbation, during the LLR - though not as early as the SLR. This matches our prediction for the task-specific hypothesis and provides further evidence that the SLR is insensitive to task constraints, as previously suggested (Kurtzer, 2014; Scott, 2012; Wolpaw, 2010). According to our results, this monosynaptic reflex loop does not seem to be influenced by a general state of stress either.

To quantify the effect of our threat-of-shock manipulation, we recorded heart-rate deceleration and the skin conductance response (SCR) on the long trials, interleaved in the experiment. We found a significant increase in the SCR for threat vs. safe trials, but the HR deceleration -that was evident in the same time window- was not differentially modulated by threat compared to safe trials. These findings suggest that threat-of-shock successfully increased activation of the sympathetic nervous system, but did not differentially affect the concurrent parasympathetic arousal. We intended to induce a freezing state in our participants, which is characterized by activation of both the sympathetic and the parasympathetic nervous system, with parasympathetic dominance (Roelofs & Dayan, 2022). However, since we did not find parasympathetic activation, our EMG findings cannot be attributed to freezing, but might be explained by increased sympathetic output alone, being better described by stress instead of threat. Indeed, studies subjecting participants to tasks that increased sympathetic nerve activation - by mental arithmetic, cold exposure and sustained gripping – also reported increased stretch reflexes (Kamibayashi et al, 2009). However, since no task-related reach was made in their experiment, it is impossible to attribute their findings to either direct spinal loop (task-unspecific) or modulatory subcortical loop (task-specific) processes.

The failure to induce differential parasympathetic activation might be explained by the chosen timings in our experimental design. In most trials (960 out of 1000), a quick motor response was required soon after trial start. This may have elicited mostly sympathetic activity, since participants prepared for rapid action and not long target anticipation. This contrasts with earlier purely perceptual studies in which there was always a long delay between trial start and target onset, requiring sustained attention and sampling of the environment (e.g. de Voogd et al. (2022), Hashemi et al. (2019) and Klaassen et al. (2024)). This sampling is mostly a parasympathetic process, in which no action is required. In the present study, most trials were focused on action execution instead of sampling and as a result a sympathetic fight-or-flight state may have dominated (Roelofs & Dayan, 2022). Future studies might focus on changing the balance between activation of the sympathetic and parasympathetic nervous system, for example by using a larger number of long trials or by creating more temporal uncertainty about target appearance in the long trials, so that continued attention and sampling will be required.

Aside from expectations regarding reflex activity, the task-unspecific and task-specific hypotheses also make different predictions regarding behavioral performance on the perceptual task. The task-specific hypothesis assumed improved perceptual processes and thus increased performance under threat. However, despite including several behavioral measures to unveil such an effect, we did not find any difference in behavior between safe and threat blocks. This is at odds with both expectations from the task-specific hypothesis and reports in previous literature (de Voogd et al., 2022; Hashemi et al., 2019; Lojowska et al., 2015). For example, we had expected more erroneous initial directions in the safe condition, but found hardly any errors and certainly no difference between conditions. However, our perceptual task was easy compared to earlier tasks, like the contrast discrimination task used by De Voogd et al. (2022), and performance effects of threat states might only become apparent in more challenging circumstances. Future work could test this, for instance by using random dot motion tasks (Adelson, 1985; Selen et al., 2012), in which the required movement direction needs to be inferred from accumulating sensory information. This paradigmatic change might at the same time also lead to parasympathetic activity, since a random dot motion task requires continuous sampling of the environment. If such a study would be successful in inducing a freezing state with both sympathetic and parasympathetic activity, the stress induced change in reflex EMG might move forward in time and/or increase in magnitude if the task-specific hypothesis holds.

## Conclusion

We conclude that stress affects motor preparation in a task-specific way. This finding suggests that earlier observed improvements in performance under threat are caused by upregulation of cortical visuomotor circuits, rather than tonic effects on peripheral reflex-loops.

## Author contributions

Yvonne Visser: conceptualization, formal analysis, investigation, methodology, software, visualisation, writing – original draft, writing – review & editing. Bob Bramson: methodology, supervision, writing – review & editing. Pieter Medendorp: conceptualization, supervision, writing – review & editing. Karin Roelofs: conceptualization, supervision, writing - review & editing. Luc Selen: conceptualization, funding acquisition, methodology, supervision, writing – review & editing.

## Data availability statement

The data for this project were collected in 2024. Preprocessed data and analysis scripts are available on the Radboud Data Repository, via the following link: https://data.ru.nl/login/reviewer-3221989340/3XBSDJ6E4UNWWSGJYP5MPBKIXHNMOTXW4YCUR2A. This study was not preregistered.

## Ethical statement

The study was performed in accordance with the declaration of Helsinki. Ethical approval was obtained from a local ethics committee (approval number ECSW-2022-144), and participants provided written informed consent before participating.

## Acknowledgements

The authors have nothing to report.

## List of abbreviations

BPM: beats per minute
ECG: electrocardiography
EMG: electromyography
GSR: Galvanic skin response
LLR: long-latency stretch reflex
M1: primary motor cortex
MT: medial temporal area
PAG: periaqueductal grey
PMd: dorsal premotor cortex
SCR: skin conductance response
SLR: short-latency stretch reflex
V1: primary visual cortex
vlPAG: ventrolateral periaqueductal grey

